# Characterization of plasma membrane proteins from maize roots (Zea mays L.) under multiple abiotic stresses *using* LC-MS/MS technique

**DOI:** 10.1101/2020.11.14.382937

**Authors:** Suphia Rafique, Nazima Nasrullah

## Abstract

The aim of this study was to understand the response of maize inbred plants showing tolerance when exposed to various abiotic stresses (drought x low-N and waterlogging x low-N stress) simultaneously. The plants under stress expressed higher photosynthetic efficiency, increase in plant height, leaf area, and were able to maintain relatively high leaf relative water content and less decrease in morphological parameters. Therefore, to understand the processes controlling the tolerance to various stresses we analyzed maize roots plasma membranes proteome of treated plants only using LC-MS/MS techniques. The large number of proteins (295) were identified which were mainly trans membrane proteins, low abundance proteins, and root specific proteins. Further, a few proteins were selected like high-affinity Nitrate transporter, NR enzyme, PEP carboxylase, and Glutamine synthetase proteins their induction were validated by qRT-PCR approach in control and treated plants. The qRT-PCR results indicated the gene of all four proteins were expressed in treated and control plants. We concluded the high-affinity nitrate transporter proteins might represent the executive part of the protective response that plays a significant role in low-N stress tolerance. The presence of other major proteins like kinases, stress-responsive TFs, calmodulin, aquaporins, stress-related proteins, and many more proteins and their interaction with nitrate transporter proteins and their role can be validated only after comparing it with control samples.

## 1. Introduction

Agricultural system is widely affected with several abiotic stresses. Recent climate prediction models indicates that rise in temperature, frequent occurrence of drought, flooding and heat waves **(IPCC, 2007 and 2008)** all are causing higher agricultural production loses. Therefore, to understand the plant response to various abiotic stresses is utmost crucial. Plants are sessile organism, their survival depends on coping with the environmental challenges. The abiotic stresses either singly or in combination causes significant damage to crop plants. The combination of two different stresses might have synergistic effect that may enhance tolerance of the plan or severe negative effect **Mittler R (2006)**. Though the plant response to different stresses is highly complex and involves changes at the transcriptome, proteome and physiological levels **Atikson and Urwin (2012)**. Perhaps, the cellular level changes in protein expression are perceived signals through the plasma membrane and are transferred to the cell. Plasma membranes are structural barrier through which exchange of substances and information are communicated to the extracellular environment of the cell **(Marmagne et al. 2004).** Most of the signal and transporter proteins are embedded in the plasma membrane these integral membrane proteins contains trans-membrane domains (hydrophobic nature), they cannot be solubilized in buffer for 2DE (**Ephritikhine et al. 2004**). Therefore, LCMS/MS-based proteomics is a parallel method with higher efficiency to identify diverse arrays and a wide range of proteins, particularly basic proteins, low-abundance, excessively large and small proteins and hydrophobic proteins **(Zhang et al. 2007**). Also it is consider highly sensitive, accurate method for identifying the integral membrane proteins. Maize is a staple food crops in the tropical climate, largely grown in marginal areas of rain fed system. during summer rainy season it has to face both drought and waterlogging stresses due to uneven distribution patterns of monsoon rains in the region (**Zaidi et al. 2008**). However at the beginning of plant growth, occurrences of these two stresses may limit the photosynthetic ability of leaves and biomass gain at the vegetative stage. Further nitrogen availability is low under both these stresses, N-uptake affected because of water deficit, while, in waterlogging leaching and de-nitrification of soil nitrogen (**Rathore et al. 1996**). Nitrogen is the major nutrient that influences the growth of plants and roots are the main organ through which mineral nutrients are taken up. Roots are the first organ that perceives abiotic stresses signals, weather due to anoxia (waterlogging), cell wall remodeling under water deficit or nutrient deficiency (N or P deficiency) henceforth they are essential for plant growth, survival and fitness. Since roots comprise a key component in the tolerance mechanisms of plants to abiotic stresses and also, it has been reported that the root is a useful tissue for proteomic research. The aim of this study was to investigate the role of root plasma membrane proteins that might be involve to sustenance of maize plants under various combined abiotic stress conditions. Therefore, physiological studies and roots proteome studies was done using LCMS/MS technique.

## 2.0 Materials and Methods

### Plant materials and stress conditions

Maize seeds were obtained from the International Maize and Wheat Improvement Center (Spanish acronym; CIMMYT®). [50-VL1018393; 51-VL0512387; 52-VL0512388; 53-VL1012838; 55-VL1018413; 56-VL0512393; 57-VL1018418; 58-VL1018419; 59-VL1018513; 60-VL1018514]. This identified inbred seeds having distinct difference in terms of tolerance/susceptibility to single stress, like waterlogging, drought and low-N. Maize seeds were sown in earthen pots (10 cm diameter) filled with sandy loam soil. The pots were kept in a naturally lit greenhouse, with air temperature 25°-30° C and relative humidity 55-65%. Ten plants (one inbred line) were chosen with six replications per plants, each pot contains 2 plants per pot after seedling emergence. Thirty pots for control and 30 for treatment, the nutrient applied in pots was calculated on the basis per kg soil, a full dose of phosphorous potash and zinc was mixed in the soil before sowing with no added organic fertilizer. (As per agronomic recommendation is N 120 kg/ha (urea) required by maize plants) However, for Low-N (LN) treatment, 25% N was used only once (195mg N/pot), unlike in control pots normal N rates (780mg N/pot) was given in split doses. The waterlogging stress was given 30 days after sowing, (**Fig 1a, b)** for up to 7 days. After completion of the stress, water was drained out from the pots by opening the holes at the bottom. Subsequently, the drought stress was given 40 days after sowing, (without recovery period) by withholding the water for a 10 days **(Fig 2a, b)**. During the stress period soil moisture content of the pots was measured on the 7^th^ day of stress from control and treatment pots **(Fig S1 Supplementary)**.

**Fig 1a, b.**
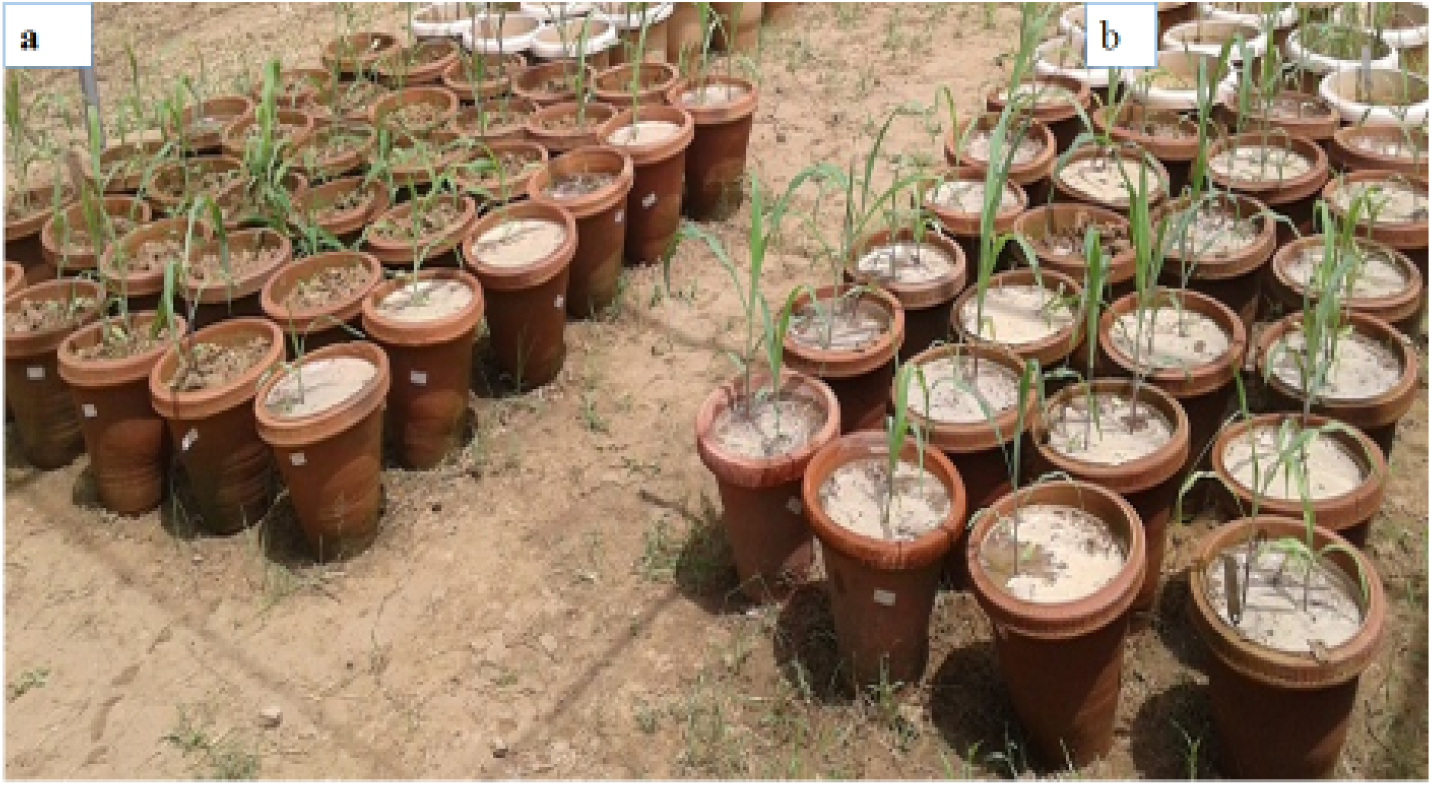
Maize plants shows no phenotypic differentiation between Control (a) and Stressed (b) plants under waterlogging x low-N stress (5 days after stress). The stress symptoms are not visible in treated plants like leaf wilting, chlorosis/necrosis, lodging and white tips on surface rooting.

**Fig 2a, b.**
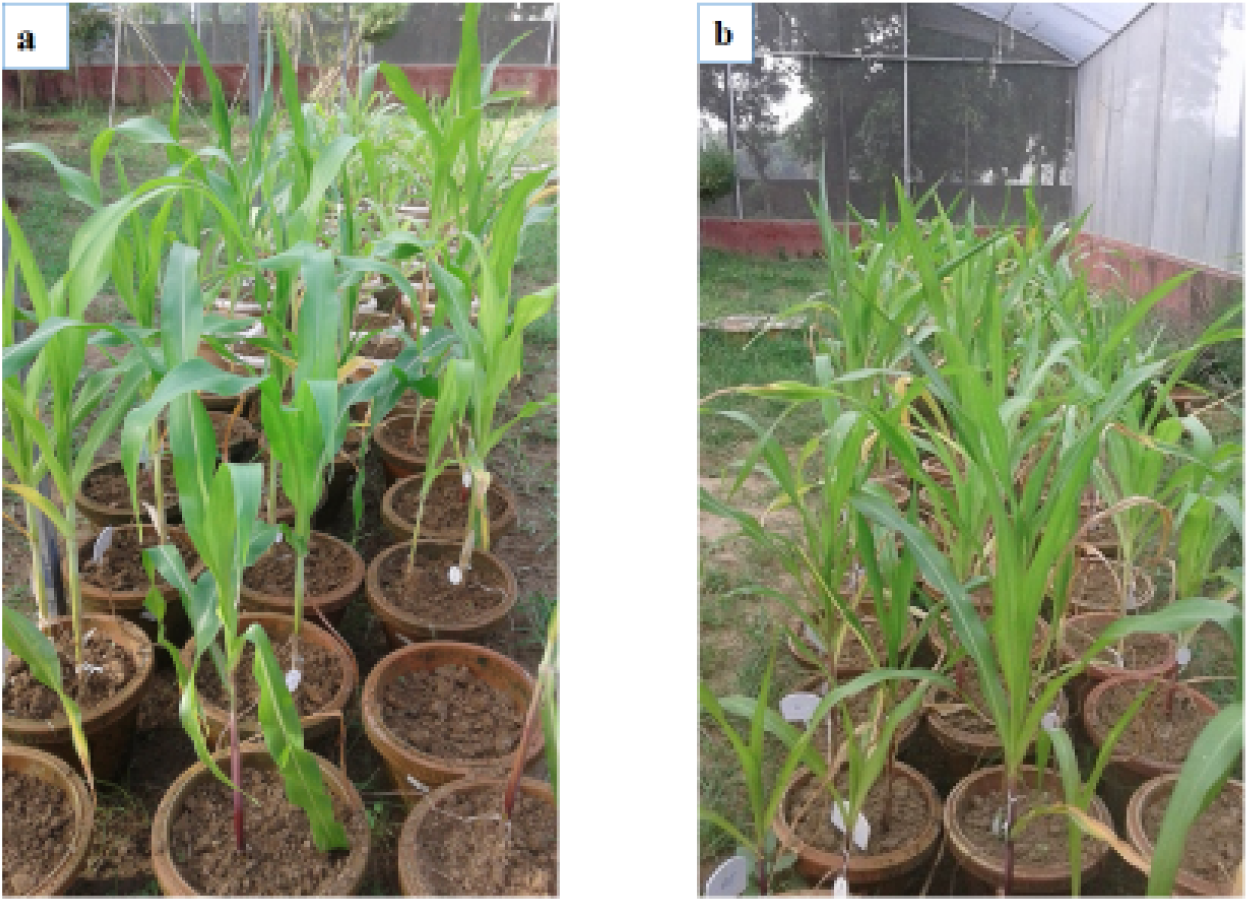
Maize plants under drought x low-N stress control (a) and stressed (b) both the phenotypes (control and stressed) are similar.

### Parameters measured under various stresses and Statistical analysis

Growth and morphological observation was recorded like, leaf area, plant height, leaf number, fresh and dry weights of shoots and roots, total fresh and total dry weights of seedlings. Physiological parameters like, Net photosynthesis rate (P_N_), stomatal conductance (gs), internal CO_2_ concentration (Ci), transpiration rate (E) was measured using LI-6400 (LI-COR Lincoln NE) portable closed gas exchange system from three plants of each control and treated plants on fully expanded leaf blades between 8:30 am to 10:00 am, also determined the chlorophyll content and leaf relative water content (RWC). For all the measured parameters each pot represented one replication. A minimum of three pots were sampled for all observation, the average of three replicates were analyzed using descriptive statistics and paired T-test for each trait and data was expressed on a per plant basis. To verify the significance of the variations of all the parameters, One-way analysis of variance (ANOVA) followed by the post hoc Tukey test (p < 0.01) was used.

### Extraction of the membrane proteins

Three biological replicates of roots sample (fresh material) were collected from the treated seedlings for proteomics and qRT-PCR studies. They were pooled and immediately frozen in liquid nitrogen and stored at −80°C for further use. However, for qRT-PCR experiment, the three different roots were collected from control plants also.

### Homogenate preparation and separation of membrane proteins

All extraction procedures were carried out on ice at 4°C. Fresh roots were weighed (5 g) in triplicate (biological) and the tissue was first grounded in liquid N2 then in 10 ml of cold extraction buffer (250 m Sucrose, 1 mM EDTA, 10 mM Tris HCl buffer, pH 7.2 and protease inhibitor) *(Sigma P9599*).Then extraction was transfer to centrifuge tube and sonicated using two 10 second pulses (30 seconds in between pulses) using a probe sonication *(Bath sonication, 30 KHz frequency*), samples kept in ice bath, to minimize the sample-air interface foaming. The intact cells, nuclei and cell debris was removed by centrifugation of the homogenate at 15000 x g for 15 minutes at 4°C (the step was repeated) and the pellet was discarded. Again the supernatant was centrifuge at 100,000 x g for 1 hour at 4°C. The obtained supernatant contains the soluble proteins that was discard. The pellet was washed by homogenization buffer and re-centrifuge at 100,000 x g, 4°C, for 1 hour. The supernatant was discard and the remaining pellet contains all of the cell’s membrane fraction was kept.

### Phenol/Ammonium Acetate-Methanol Precipitation of membrane proteins

Membrane pellet was suspend in 0.5 ml of extraction buffer (0.7 M sucrose, 0.5 M Tris, 30 mM HCl, 50 mM EDTA, 0.1 M KCl, 2% (v/v) 2-mercaptoethanol and 2 mM PMSF) (*Sigma-Aldrich*) and homogenize. Incubated for 10 min at 4°C and then an equal volume of Tris-saturated (7.5 pH) phenol was added. Centrifuged to separate the phases, the phenol phase was recovered and re-extract with an equal volume of extraction buffer. Proteins was precipitated from the phenol phase by adding of 5 volumes of 0.1 M ammonium acetate in methanol and incubated at −20°C for overnight. The precipitate was washed 3-times with ammonium acetate in methanol and one time with acetone, the pellet was air dried and solubilize in rehydration buffer by incubating for at least 1 hour at room temperature, with occasional vortex and then centrifuge at 100,000 x g, 4°C, for 1 hour. The supernatant was removed and save. The sample protein concentration were quantified by Bradford’s method. **Bradford (1976)** using bovine serum albumin (*Fischer Scientific*) as a standard.

### Trypsin digestion in solution samples and data analysis in LC MS/MS

The goal of this study was to comprehensive identification of integral membrane proteins. 100μg of roots protein sample was taken for digestion; the volume was made up to 100μl with 50mM NH4HCO3. The sample was treated with 100mM DTT at 95°C for 1 hour, followed by 250mM Iodoacidamid (IA) at room temperature in the dark for 45 min. The sample was then digested with trypsin and incubated overnight at 37°C. The sample was vacuum dried and dissolved in 10μl of 0.1% formic acid in water. After centrifugation at 10000 x gs, the supernatant was collected into the separate tube. 1μl injection volume was used on C18 nUPLC column for separation of peptides, and then followed by analysis on the *Water Synapt G2 Q-TOF* instrument for MS and MSMS. The raw data was processed by *MassLynx 4.1 WATERS*. The individual peptides MS/MS spectra were matched to the database sequence for proteins identification *on PLGS software, WATERS*. For Protein identification Database used UNIPROT, Mass Tolerance; 50ppm and Peptide mass tolerance; 100ppm for the search to proceed. Specific modification; Carbamidomethyl and variable modification; Oxidation (M). Result based on the Scores of the matching protein masses and probable peptides was given as output.

### Quantification of gene expression by Semi-quantitative RT-PCR

Total RNA was extracted from 250 mg of frozen roots tissue stored at −80° C of treated (after combined stress treatment) and control plants, using the TRIzol method (*as described by the manufacturer*) An aliquot of total RNA was treated with RQ1 RNase-free DNAse (*Promega*), to avoid genomic contamination and 1μl of total RNA was quantified by spectrophotometer using a Nanodrop 1000 (*Thermo Scientific, Nanodrop Products*). First cDNA was synthesized from 100 ng of total RNA and mixed with 1 μl of Oligo dT (10 μM). The reaction was incubated 5 min at 70°C, qRT-PCR was performed with gene-specific primers corresponding to the genes encoding the identified proteins. **Table S1 (Supplementary)**. The primers were designed to generate PCR products of 500-1000bp. MEP and LUG used as reference gene for normalization of internal cDNA input.

## 4.0 Results and Discussion

### Effects of various stresses on maize plants

This studies was carried out with the purpose to understand the effects of various stresses applied simultaneously on maize inbred plants at vegetative stage and to reveal the process of tolerance by proteomic approach, specifically the low-N response. We have subjected the maize plants to various abiotic stress in combination like waterlogging x low-N and drought x low-N and observed the photosynthesis rate, stomatal conductance (gs) and intercellular CO_2_ (ci) in treated plants **(Fig 3a, b).**

**Fig 3a and b.**
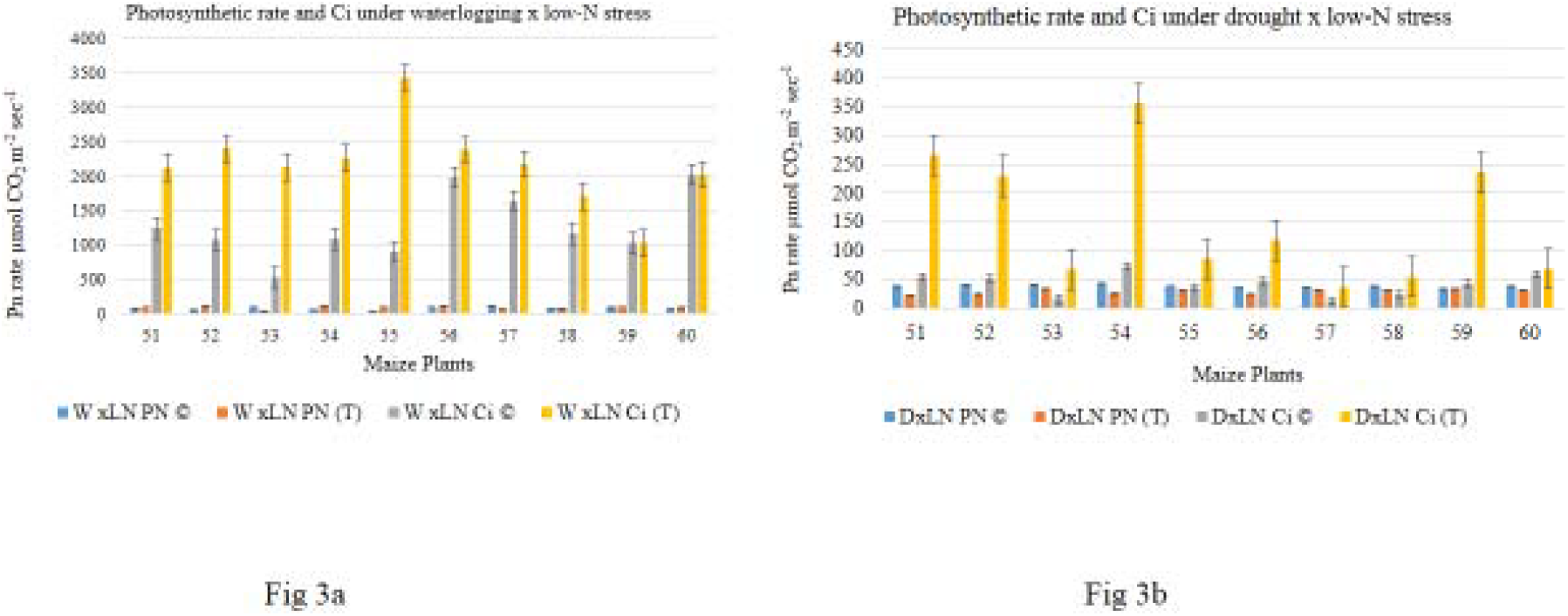
Photosynthetic rate of control and stressed plants under waterlogging x low-N drought x low-N stresses. Mean values. of Pn rate and internal CO_2_ concentration (Ci) are significant at p < 0.05

We found that under waterlogging x low-N stress, all treated plants shows higher photosynthetic efficiency (13.77%) relative to control plants. Whereas, in drought x low-N stress the decline in photosynthetic rate was observed compared to control plants, though the decline in photosynthesis was not below the range 40-50μmol CO_2_ sec^−2^mol^−2^ under drought. Photosynthesis is among the primary process to be affected under stress condition. However, under multiple stress conditions decline in photosynthesis may be due to oxidative stresses **(Chaves and Oliveira, 2004),** these multiple stresses affect leaf photosynthetic machinery (**Ort, 2001**). Similarly, **Zaltev and Lidon (2012)** indicated the sustenance of photosynthetic mechanism by plants under water deficit stress shows the drought tolerance capability. Hence our results indicated that overall photosynthesis sustained in treated plants under multiple stresses. In our experiments plants were grown under low-N stress yet maintain photosynthesis. Although photosynthetic rate and leaf N had the strongest correlation to AEI (assimilation efficiency index) in maize, **Settimi and Maranville (2008)**. The phenotypic observations of treated plants under waterlogging x low-N stress shows no lodging, wilting, leaf necrosis, and surface rooting, instead early brace root development was observed. Subsequently in drought x low-N stress symptoms includes, leaves drooping, yellowing, wilting and premature leaf there was no such phenotype was visible in treated plants. Also growth, morphological, and physiological parameters have non-significant difference in the mean values between control and treated plants **(Table 1**).

**Table 1.** The growth, morphological and physiological traits measured under various abiotic stress combination in control and stressed maize plants. Mean ± SE (m) and C.D values are shown in table.

| Trait | Treatment | Waterlogging x Low-N stress |  |  | Drought x Low-N stress |  |  |
| --- | --- | --- | --- | --- | --- | --- | --- |
|  |  | Mean | C.D | SE(m) | Mean | C.D | SE(m) |
| Photosynthesis rate ($\mu\text{mol CO}_2 \text{ m}^{-2} \text{ S}^{-1}$) | Control | 80.733** | 24.1 | 8.401 | 38.233* | 6.35 | 2.21 |
|  | Treated | 93.68 |  |  | 28.623 |  |  |
| Conductance (gs) | Control | 0.087** | 0.023 | 0.008 | 0.264** | 0.107 | 0.037 |
|  | Treated | 0.101 |  |  | 0.142 |  |  |
| Internal $\text{CO}_2$ (Ci) | Control | 1819.47* | 1785.61 | 622.433 | 57.267** | 117.098 | 40.818 |
|  | Treated | 1902.20 |  |  | 136.315 |  |  |
| Transpiration rate E (mol) | Control | 2.428** | 0.943 | 0.329 | 4.742** | N/A | 0.538 |
|  | Treated | 2.722 |  |  | 2.83 |  |  |
| Plant height (cm) | Control | 33.9** | N/A | 3.251 | 29.167** | N/A | 3.234 |
|  | Treated | 31.5 |  |  | 28.933 |  |  |
| Leaf area ($\text{cm}^2$) | Control | 1746.54** | N/A | 334.343 | 1344.42* | N/A | 321.882 |
|  | Treated | 1146.85 |  |  | 1236.78 |  |  |
| Leaf No. (per plant) | Control | 7.8* | N/A | 0.699 | 6.633* | N/A | 0.587 |
|  | Treated | 7.0 |  |  | 6.667 |  |  |
| Fresh shoot weight (g) | Control |  |  |  | 72.42** | N/A | 12.9 |
|  | Treatment |  |  |  | 45.215 |  |  |
| Dry shoot weight (g) | Control |  |  |  | 14.718* | N/A | 3.094 |
|  | Treatment |  |  |  | 9.857 |  |  |
| Fresh root weight (mg) | Control |  |  |  | 10.12** | 7.061 | 2.461 |
|  | Treated |  |  |  | 4.243 |  |  |
| Dry root weight (mg) | Control |  |  |  | 2.43** | N/A | 0.613 |
|  | Treated |  |  |  | 1.042 |  |  |
| Total fresh weight (g/plant) | Control |  |  |  | 85.643* | N/A | 17.887 |
|  | Treated |  |  |  | 51.215 |  |  |
| Total dry weight (g/plant) | Control |  |  |  | 17.154** | N/A | 3.481 |
|  | Treated |  |  |  | 11.442 |  |  |

\*\*significant difference for $p < 0/ 0.001$
\*significant difference for $p < 0.05/ 0.01$

Therefore, it appears that might be some changes during the stress or some signals were regulated to overcome the stressful conditions, although in present work plants were subjected to various stresses yet they maintain high assimilation rates. Therefore to determine whether the observed rates of photosynthesis described above correlated with changes in proteins under stress conditions. The complete protein profile of treated plants was analyzed in detail by LCMS/MS method (in solution).

### Characterization of proteins identified by LC-MS/MS technique (in solution)

The aim of the study was to understand the response of maize plants under multiple stress conditions. Maize plants subjected to combined stresses (waterlogging x low-N and drought x low-N) and the proteins were extracted from roots and analyzed by LC-MS/MS technique (in solution). In the root tissue of maize, approximately 295 proteins were identified shown in **Table 3**.

**Table 3:** Plasma membrane proteins of maize roots expressed under combined abiotic stresses and identified by LC-MS/MS technique

| Protein Name <sup>a</sup> | Accession No <sup>b</sup> | mW(Da) <sup>c</sup> | pI(pH) <sup>d</sup> | Peptide <sup>e</sup> | Theoretical <sup>f</sup> | %Coverage <sup>g</sup> |
| --- | --- | --- | --- | --- | --- | --- |
| **CALCIUM BINDING PROTEINS** |  |  |  |  |  |  |
| Calmodulin (fragment) | U5Q018_MAIZE | 12554 | 4.04 | 24 | 15 | 100 |
| Calmodulin (fragment) | U5Q0B5_MAIZE | 11879 | 4.08 | 21 | 13 | 97 |
| Calmodulin (fragment) | U5PZT9_MAIZE | 10511 | 4.16 | 21 | 11 | 100 |
| Calmodulin (fragment) | U5Q0D5_MAIZE | 9305 | 4.07 | 16 | 10 | 100 |
| Calmodulin (fragment) | U5Q0C3_MAIZE | 12291 | 3.83 | 19 | 12 | 100 |
| Calmodulin (fragment) | U5PZT0_MAIZE | 12703 | 4.01 | 20 | 14 | 98 |
| Calmodulin (fragment) | U5Q0C8_MAIZE | 10553 | 4.16 | 17 | 11 | 96 |
| Calmodulin (fragment) | U5PZQ0_MAIZE | 10665 | 4.19 | 23 | 11 | 95 |
| Calmodulin (fragment) | U5Q3A9_MAIZE | 12583 | 4.04 | 26 | 16 | 100 |
| Calmodulin-binding protein | Q41796_MAIZE | 14604 | 5.06 | 21 | 8 | 69 |
| Calcium-binding protein | Q43712_MAIZE | 47983 | 4.28 | 46 | 45 | 91 |
| Calcium transporting ATPase | A0A096PNP4_MAIZE | 97157 | 7.59 | 107 | 56 | 87 |

Most of the proteins were related to diverse biological functions and has been categorized according to **Bevan et al. (1998).** The protein percentage was calculated by dividing the type and number of functional proteins from total no proteins present. Proteins related to nitrogen, and carbon metabolism were maximum in number (44.6%). However, some metabolism related proteins were associated with plasma membranes like, Enolase, G-3P dehydrogenase, PEP carboxylase, Nitrate Reductase enzyme. But some of the enzymes are soluble proteins, **Alexandersson et al. (2004)** consider them as contaminants of the plasma membrane preparation. The second maximum were uncharacterized proteins (19%), calmodulin, kinases (Signal transduction), transcription factors (TF), transporter proteins, root specific proteins, cell division, translation and cell wall synthesis proteins, stress related proteins. While others were hormones, ubiquitin related proteins and some proteins with unknown functions shown in **Fig 4**.

**Fig 4.**
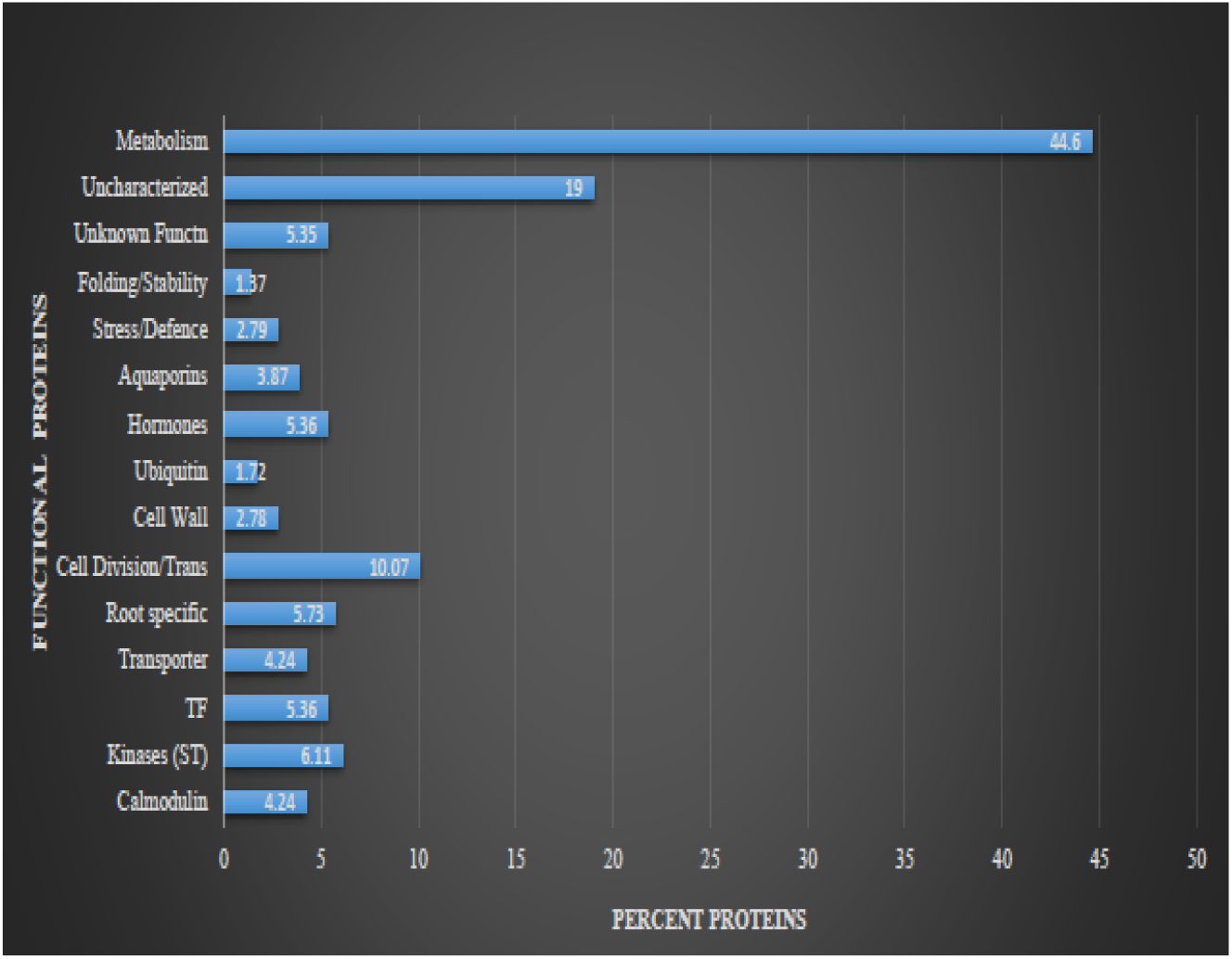
Functional classification of plasma membrane proteins of maize root identified by LC-MS/MS technique. Functional classification of expressed proteins under combined stresses in plasma membranes of maize roots. The numbers and percentages of proteins from each functional category assigned on the basis of identified proteins. Proteins were categorized using the criteria of Bevan et al. (1998).

The concentration or abundance molecules in the sample can be detected by mass spectrometry. In our experiment, the maximum percentage coverage 95.41%, 94.58% and 75.38% is shown by NRT2.1 protein (Accession No’s: Q53CL7, Q0VH26, and Q0VH25). The highest peak of the chromatogram **Fig S3 (supplementary)**, also indicate the high expression of NRT2.1 proteins in the roots samples. Another maximum percent coverage was membrane bound Nitrate Reductase proteins (Accession No. Q4U5G4, 95.35%).

### Possible role of NRT2.1 proteins, Nitrate Reductase, Phosphoenol pyruvate carboxylase and Glutamine synthetase and their validation using qRT-PCR studies in control and stressed plants under multiple stresses

The maize plants were grown under low-N stress, and we have identified the nitrate transporter protein in treated plants. The mRNA transcript of high affinity nitrate transporter in treated plants was spotted by qRT-PCR studies, **Miller et al. (2007)** emphasize the regulation of HATS at the range of above 1mM concentrations (Soil low-N), had played important role in sensing NO_2_ availability at root surface through feedback regulation of expression and phosphorylation of proteins. Thus confirms their induction at low NO^−^_3_ concentration. Similarly, the induction of NRT2.1 genes at very low levels of NO^−^_3_ (10–50 mM) was noted in *N*. *plumbaginifolia* and *Arabidopsis* **(Krappet al. 1998; Filleur and Daniel Vedele 1999)**. The nitrate transporter transcripts was also detected in control plants **Fig 5**.

**Fig 5.**
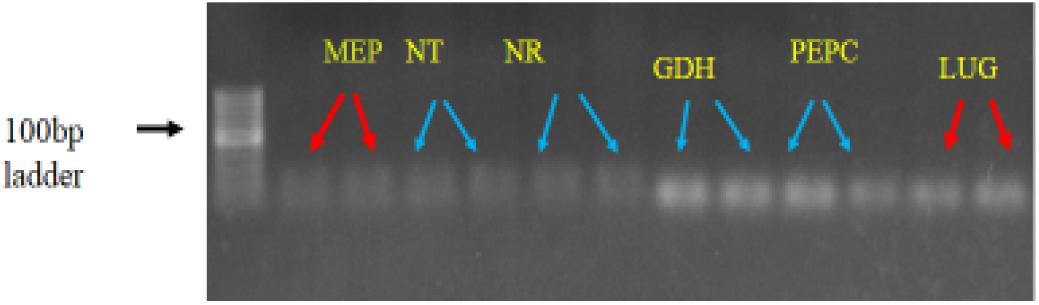
Gene expression analysis by qRT-PCR of 4-plasma membrane proteins. Transcripts of four plasma membrane proteins shows the induction of all proteins in control and stressed plants. Represented with blue arrow- Nitrate transporter (NT), Nitrate Reductase (NR), Gulamine synthetase (GDH) and Phosphoenol pynivate carboxylase (PEPC) RED arrow-MEP and LUG internal control genes of Maize.

The presence of two types HATS was detected in barley the one was iHATS, and other cHATS, at low NO^−^_3_ concentration **(Siddiqi et al. 1990)**, similar to our results the constitutive HATS (cHATS) in control plants whereas, inducible HATS (iHATS) transcripts in treated plants. Moreover, possible role of NRT.2 proteins might to maintain N homeostasis in various stresses. In similar work the roots of two salt cultivars (salt-tolerant FL478 and salt-sensitive IR29 rice varieties) had shown significant up regulation of gene encoding nitrogen transporter in both the cultivars. Up-regulation of nitrogen transporter gene maintain the N homeostasis in tolerant cultivars, whereas, in salt sensitive cultivars up-regulated gene significantly reducing the N content **(Senadheera et al. 2009).** Therefore transporter gene may have contributed to the salinity tolerant in both the cultivars. Another study shows higher abundance of transcripts related to high affinity nitrate transporters (NRT2.2, NRT2.3, NRT2.5, and NRT2.6) in tolerant genotype of barley **Gelli et al. (2014)**. However, soon after sensing NO^−^_3_ concentration in external medium plants respond by activating genes encoding NO^−^_3_ transport system and many enzyme systems **Gojon et al. (2011).** In barley plants, Nitrate Reductase is the first enzyme to be involved in assimilation (**Crawford 1995; Wang et al. 2004**; **Ho et al. 2009 Krouk et al. 2010).** Successively, strong correlation between increased rates of NO^−^_3_ uptake and NR activity was observed **(Larsson and Ingemarson 1989; Jackson et al. 1986).** The presence of NR transcripts in treated plant correlates with the results of proteomics, that second highest coverage of protein was membrane bound NR in stress plants. The transcripts of glutamine synthetase were correspondingly expressed in treated genotypes. Consequently it shows amino acid synthesis was in progress in treated plant. Likewise, **Li et al. (1993**), detected the presences of genes of GS1-1 form and its expression in roots and confirms, the assimilation of NH_4_^+^ by the glutamine synthetase pathway for the amino acid synthesis. Moreover, in treated plants transcripts of PEPC (Phosphoenolpyruvate carboxylase) expressed. A study showed the exogenous supply of nitrogen selectively increased the levels of protein and mRNA for PEPC, in parallel, glutamine synthetase activity and/or ferredoxin-dependent glutamate synthase increases. Also with the steady-state level of PEPC mRNA and the major amino acids, glutamine level increased for 7 hours after nitrogen supply **(B Sugiharto and T Sugiyama, 1992)**. The presence of these proteins correlates to carbon metabolism, on incubating of maize roots for 30h in nitrogen nutrition, enhances the enzymes involved in nitrogen metabolism and carbon metabolism showed by **Prinsi et al. (2009).** However, **Nohzadeh et al (2007)** performed real-time PCR analysis, to investigate the correlation between mRNA and protein levels, in the roots of PM of salt-tolerant variety of rice, IR651, for three salt responsive genes (1,4-benzoquinone reductase, a putative remorin protein, and a hyper sensitive induced response protein). In their results no correlation was detected between the changes in the levels of gene and protein expression. Whereas, our results show correlation between plasma membrane proteins and mRNA level. Though in our study plants were under low-N stress and deficient in nitrogen nutrition but due to the induction of nitrate transporter proteins and other 3-proteins, there is coordinated regulation of interaction of carbon and nitrogen metabolism. Subsequently maintains the homeostasis in various stresses.

### Gravy Index of Plasma membrane proteins

The GRAVY is a computational program that evaluate the hydrophilic and hydrophobic properties of proteins along its amino acid sequence. The GRAVY score **(Kyte and Doolittle, 1982)** takes into account the size and the charge of the whole protein and ranges. The GRAVY of the maize roots plasma membrane proteins analyzed ranges from –1.32 to 0.402. Whereas, positive values referring to hydrophobic proteins. In our roots sample highly hydrophobic proteins is plasma membrane intrinsic protein (Q84RL8_MAIZE) it’s Gravy score +0.4 The **Table 4,** shows the Gravy index of roots plasma membrane proteins along with pI and cellular location of the proteins.

**Table 4.**
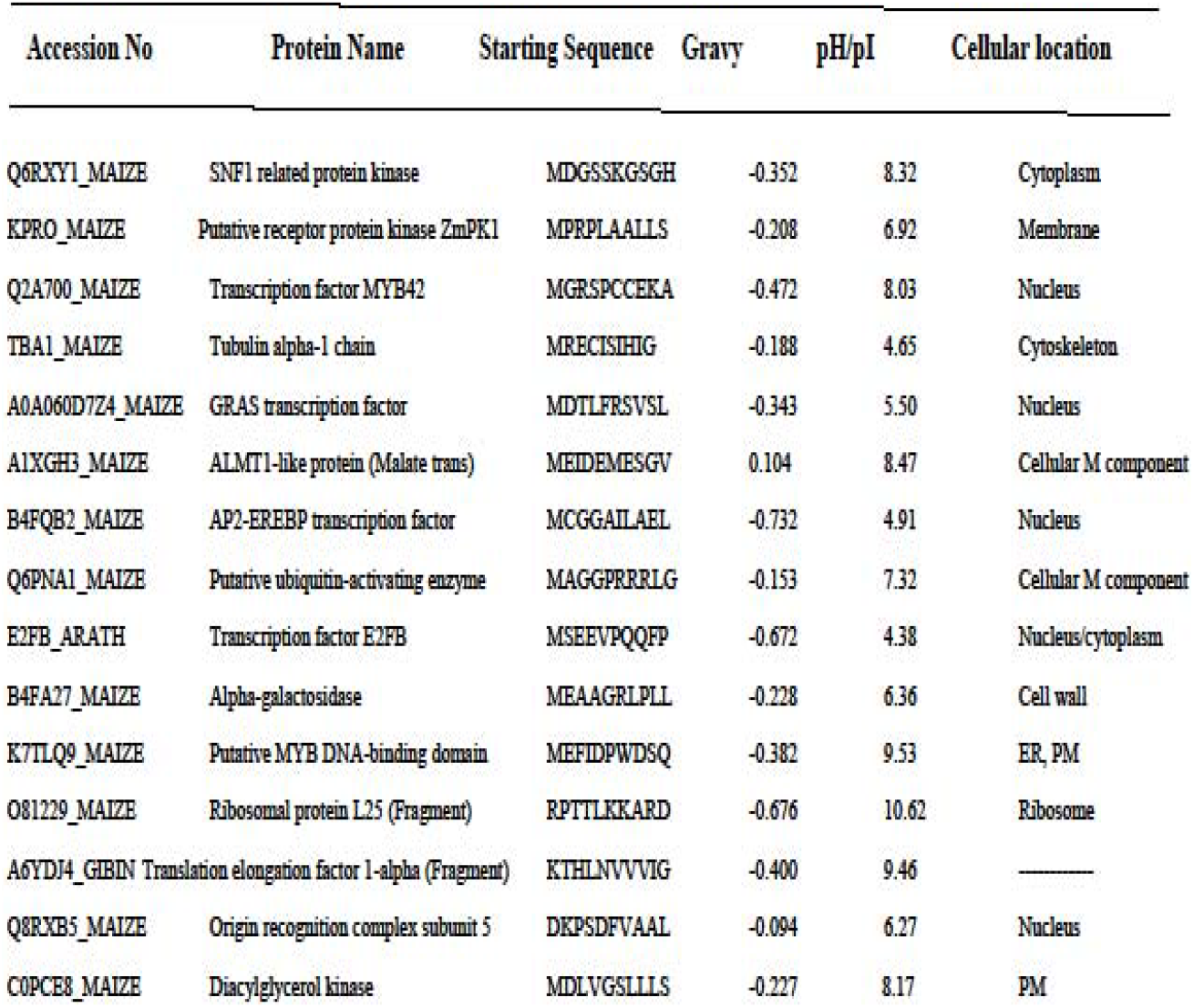
Gravy index of plasma membrane proteins of maize

### Proposed Model

Therefore, on the basis of our proteomics data, qRT-PCR studies, and physiological status of the treated plants in response to various stresses. A model has been proposed that shows the plants response in low-N stress and their combine adversity stress adaptation strategies. The induction of Nitrate transporter proteins that enhanced and involve network of proteins to maintain homeostasis in other two stresses (waterlogging and drought) thus help to acclimatize the maize plants in various abiotic stress conditions has been shown in **Fig 6.**

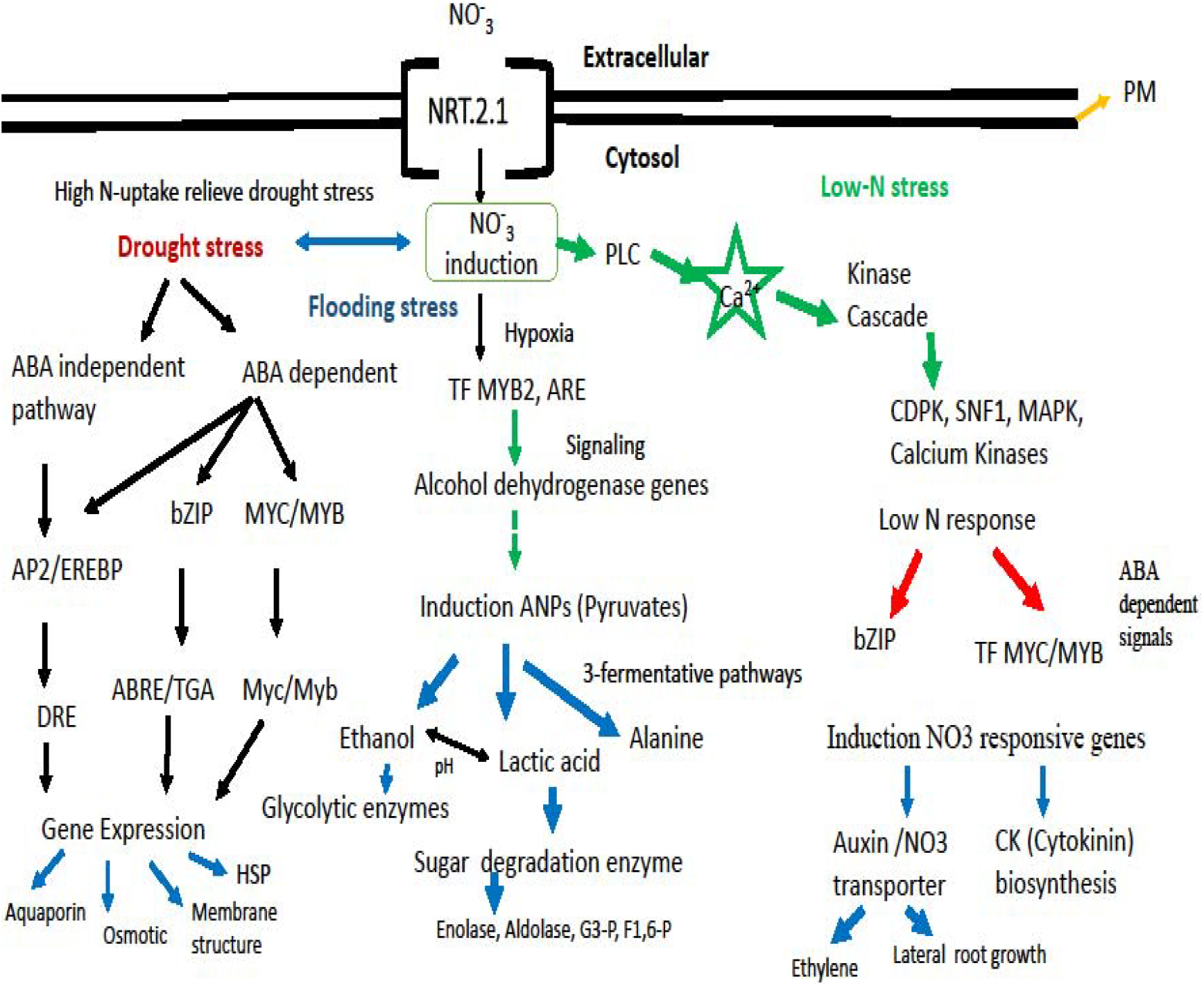

## 5.0 Conclusion

The maize plants when subjected to various abiotic stresses in combination shows tolerance. The phenotypic observations in treated plants does not exhibited any stress related symptoms. The mean values of various growth, morphological and physiological parameters in treated and control shows no difference. To understand the tolerance mechanism under multiple abiotic stresses, a study of roots plasma membrane has been completed. We have used the LC-MS/MS techniques (in solution) to identify and characterized the roots plasma membrane proteins in treated plants only. The presence of a large number of integral hydrophilic, hydrophobic and low abundance of proteins were identified in our results. The physiological analysis, and transcriptional studies validates the role of four proteins (treated plants). Further, the role of other proteins can be validated only after comparing the control roots protein samples with the treated roots proteins. In present context we can assume the role of characterized proteins of stressed plants, might be due to coordinated regulation and expression of various proteins along with induction of ‘High-Affinity nitrate transporter proteins". These are involved in sensing and transporting nitrogen in low-N condition, and trigger signaling cascades which in turn activates membrane bound TFs, that initiate transcription of genes for low-N stress, waterlogging and drought stresses. Thus, maintain metabolic homeostasis and counteracting the effects of stress and ameliorate to acclimatize the plants at vegetative stage. Similarly, **Scheible et al. (2004)** observed the direct and indirect consequences of the nitrogen availability on the whole plant metabolism.

## Supporting information

Results

