## Supplementary material for "Characterization of plasma membrane proteins from maize roots (Zea mays L.) under multiple abiotic stresses *using* LC-MS/MS technique": Materials and Methods

### Fig S1: Soil moisture content of pots in drought x low-N stress.

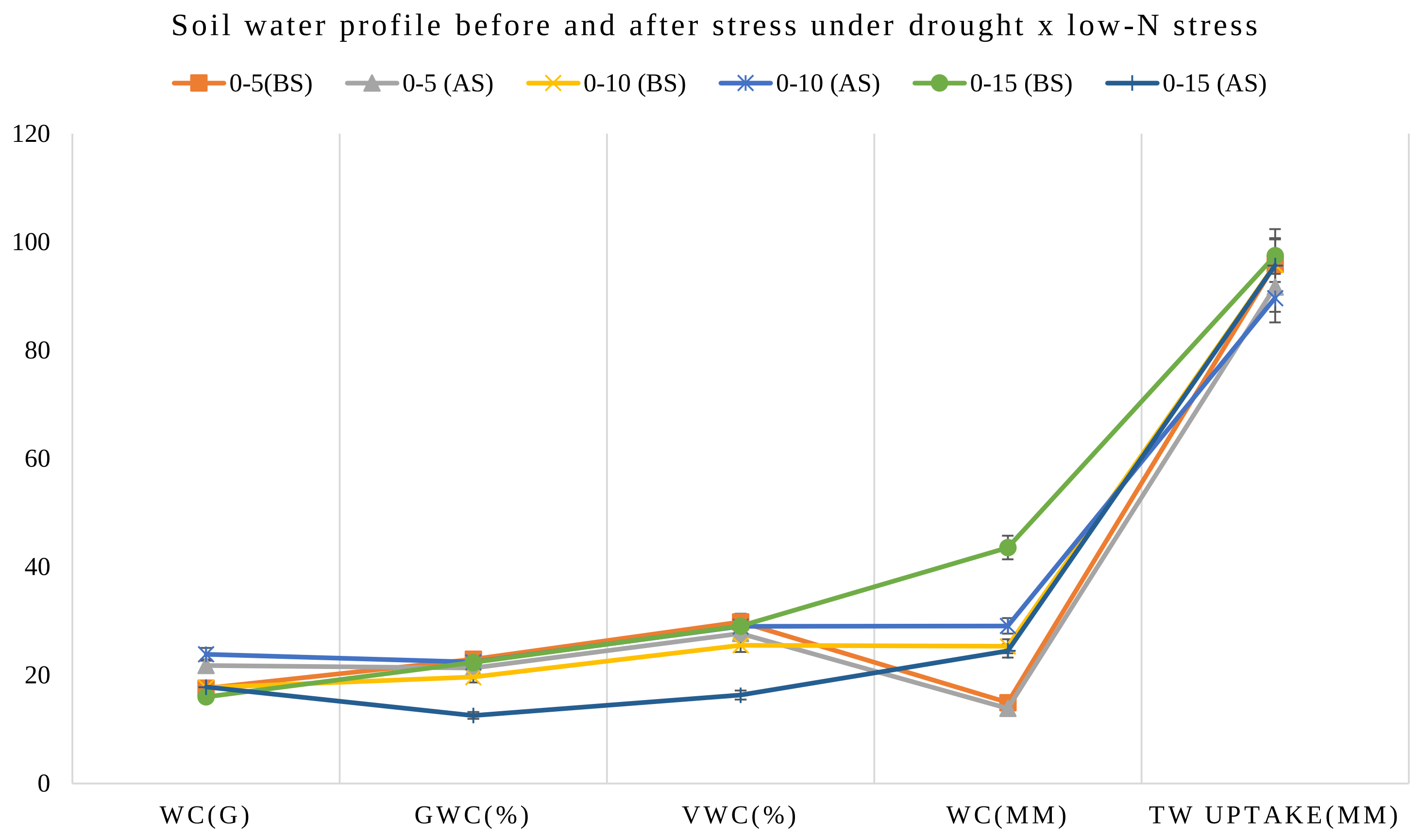


Fig showing mean values of soil moisture content in some selected pots during the screening in drought x low-N. Before stress (BS) and After stress (AS) taken at various soil depth (cm). The values are for 3-independent replicates taken in triplicate (n=9).
