## Supplementary material for "Characterization of plasma membrane proteins from maize roots (Zea mays L.) under multiple abiotic stresses *using* LC-MS/MS technique": Materials and Methods

**Table S1: Pimers sequence of 4-genes and maize reference gene for qRT-PCR experiment**

| Gene name | Primer sequence (forward/Reverse) |
|---|---|
| 1 High Affinity Nitrate Transporter (NT) | (FP) CCTCTCCTGGATCTCCTTCTTC<br>(RP) CTTGGTGAGGTTGAGGTTGT |
| 2 Nitrate reductase (NR) | (FP) GAGCTGAACATAAACTCGGTGATA<br>(RP) GGCGTATCCTTTCATGGTGTAG |
| 3 Phosphoenol pyruvate Carboxylase (PEPC) | (FP) GATTGCTTTGGCGCGTATATC<br>(RP) ACTCTCAGTGGCTGCTTTAC |
| 4 Glutamate dehydrogenase (GDH) | (FP) CTCTGAGCTTGAGAGGCTTAC<br>(RP) GTGAGTTGGTGCCCATATCT |
| 5 Housekeeping gene (MEP) | (FP) TGTACTCGGCAATGCTCTTG<br>(RP) TTTGATGCTCCAGGCTTACC |
