## Supplementary figures and images for "Characterization of plasma membrane proteins from maize roots (Zea mays L.) under multiple abiotic stresses *using* LC-MS/MS technique"

### Results

**Fig S3:** Showing chromatogram and the highest peak of NRT2.1 proteins

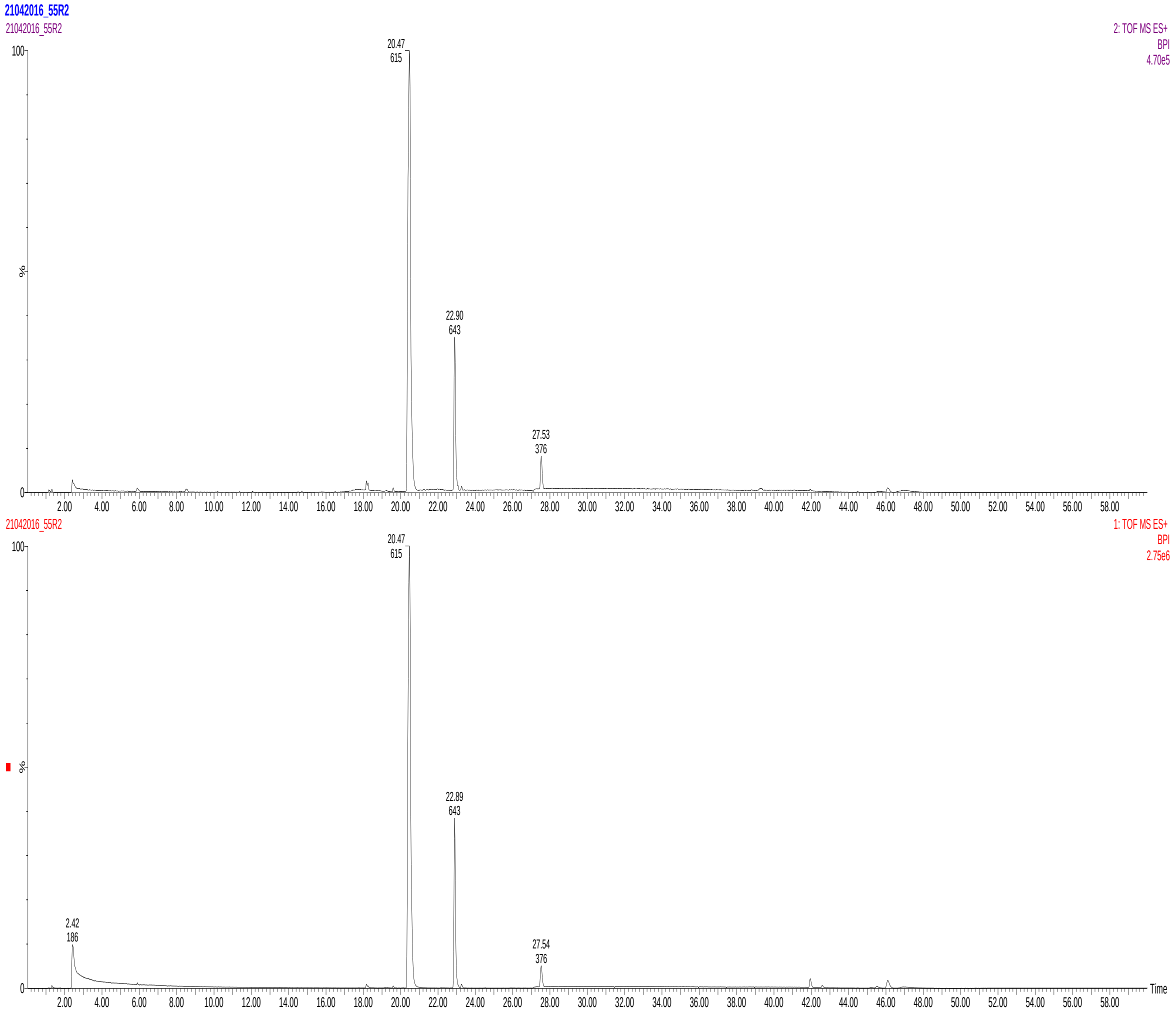
